# A structure-based tool to interpret the significance of kinase mutations in clinical next generation sequencing in cancer

**DOI:** 10.1101/2025.03.13.643138

**Authors:** Amith Rangarajan, Ilona Sviezhentseva, Emma Gunderson, Yana Pikman, Matthew P. Jacobson, Beth Apsel Winger

## Abstract

**Introduction:** Clinical workflows to analyze variants of unknown significance (VUSs) found in clinical next generation sequencing (NGS) are labor intensive, requiring manual analysis of published data for each variant. There is a strong need for tools and resources that provide a consistent way to analyze variants. With the explosion of clinical NGS data and the concurrent availability of protein structures through the Protein Data Bank and protein models through programs such as AlphaFold, there exists an unprecedented opportunity to use structural information to help standardize NGS analysis with the overall goal of advancing personalized cancer therapy.

**Methods:** Using the Catalogue of Somatic Mutations in Cancer (COSMIC), the largest curated database of clinical cancer mutations, we mapped thousands of missense mutations in the kinase and juxtamembrane (JM) domains of 48 receptor tyrosine kinases (RTKs) onto structurally aligned kinase structures, then clustered known activating mutations along with VUSs based on proximity in three-dimensional structure. Using cell-based models we illustrate that our resource can be used to identify activating mutations and provide insight into mechanisms of kinase activation and regulation.

**Results:** We provide a database of structurally aligned and functionally annotated mutations that can be used as a tool to evaluate kinase VUSs based on their structural alignment with known activating mutations. The tool can be accessed through a user-friendly website in which one can input a kinase mutation of interest, and the system will output a list of structurally analogous mutations in other kinases, as well as their functional annotations.

**Discussion:** We expect our database to be an important addition to the current tools and resources used to analyze clinical NGS, with important clinical implications to guide recommendations for personalized cancer therapy.

## Introduction

Protein kinases comprise the most targetable class of oncogenic proteins, with at least 80 FDA-approved kinase inhibitors available (1). Extensive precision medicine efforts using clinical next generation sequencing (NGS) have expanded the use of these inhibitors to virtually all cancer types (2). However, most mutations detected through NGS are variants of unknown significance (VUSs), which are genetic mutations that have an unknown effect on human health (3, 4). Kinase VUSs create a particular challenge for oncologists who have to decide whether to use kinase inhibitors for uncharacterized mutations in patients with limited treatment options (2).

Over 100 variant prediction algorithms have been created to address the challenge of VUS interpretation in NGS (5). Such algorithms, some of which are specific to disease or protein class, generally aim to predict the functional significance of mutations by incorporating multiple elements into a score and/or using machine learning/artificial intelligence (5-9). Most utilize protein sequence data, such as the degree of chemical change and evolutionary conservation at the mutated amino acid site; some also incorporate aspects of protein structure (10-13). However, the specificity of available algorithms remains at only ∼60-80%, leading the American Association of Pathology to recommend against applying them to clinical care (9, 14, 15). Clinical workflows to analyze NGS continue to be labor intensive, requiring in depth analysis of published data by molecular pathologists and other content experts who often come together as a Molecular Tumor Board (8, 16, 17). There is a strong need for clinical and experimental data across variants that could help standardize NGS analysis.

Thousands of experimental structures are now available for human protein kinases, and these experimental structures are supplemented by models generated by powerful structure prediction algorithms such as AlphaFold (18). These advances have stimulated discussion of how protein structure can advance clinical practice (19). The functional impact of a given mutation, if any, depends sensitively on its location within the three-dimensional structure of the protein. For example, a mutation that changes the net charge of the protein (e.g., Lys->Glu) might have no significant impact on function if the amino acid is located on a surface loop distant from the active site but could cause the protein to become unfolded if the amino acid is buried in the core or could cause loss of function if the amino acid is located in the active site. Thus, the effect of a mutation on three dimensional protein structure can provide important insights into the functional impact of mutations (17, 20, 21). Incorporating protein structure into mutational analysis is most effective if there is a clear understanding of how mutated protein regions contribute to protein function, as is the case for kinases. Since kinase structure-function relationships are well defined, understanding the impact of kinase mutations on protein structure can be particularly useful in providing functional insights such as whether a mutation is activating (22).

Receptor tyrosine kinases (RTKs) constitute an important family of oncogenic kinases. They are frequently mutated to drive cancer growth and they are targets for the majority of kinase inhibitors (23). RTKs have four main domains: an extracellular ligand-binding domain, a transmembrane domain, a juxtamembrane (JM) domain, and a catalytic kinase domain (Figure 1A,B). They exist in equilibrium between active and inactive states (23, 24). Active and inactive states are structurally distinct, with key conserved motifs rearranging in the active state to position catalytic residues in a geometry that enables catalysis (Figure 1B) (22, 24). For example, the activation loop, which provides a platform for substrate binding, blocks the active site in the inactive conformation and extends out of the active site in the active conformation. Also during activation, a motif called the αC-helix shifts inward to create a salt bridge that coordinates ATP for catalysis (Figure S1). For a subset of kinases (e.g. KIT, PDGFRA, FLT3), the JM domain is autoinhibitory, blocking the active site in the inactive conformation and moving away from the active site in the active conformation (25-27).

**Figure 1.**
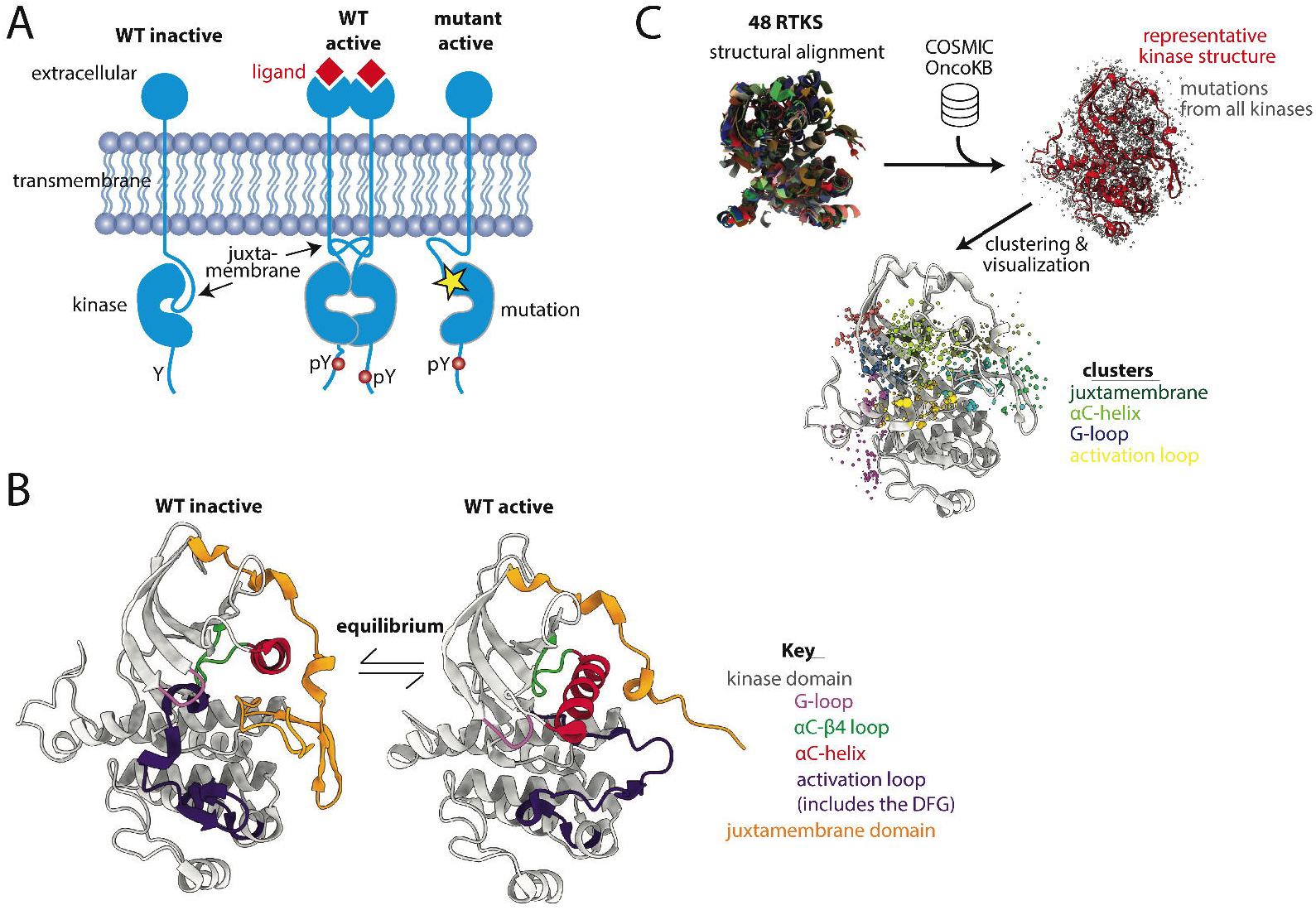
Well-defined structure-function relationships in receptor tyrosine kinases (RTKs) informed the approach to align and cluster clinical RTK mutations. A) Schematic demonstrating both endogenous (wildtype = WT) and oncogenic activation of RTKs (yellow star = mutation, pY is a phosphorylated Tyr, indicating an activated kinase). B) Key structural motifs that change conformation between active and inactive kinases. C) Basic methodology for structural alignment of mutations and clustering. For clarity, only one kinase structure is shown after the initial multi-structure alignment; grey dots are mutations, colored dots are clusters of different mutations, with each group of different colored dots comprising a single cluster. Dots do not all sit on the backbone because only a single representative structure is shown and because the mutations are on sidechains, which are not depicted, for clarity. COSMIC is the mutational database used, OncoKB is the annotation database used.

The equilibrium between inactive and active states is regulated by ligand binding and post-translational modifications, and can also be modulated by oncogenic mutations that push the equilibrium of RTKs toward the active state through mechanisms that are sometimes, but not always, known (Figure 1A, B) (24). For example, in lung cancer, short in-frame deletions in the β3-αC loop of EGFR pull the αC-helix toward the active position, leading to kinase activation (28). In contrast, KIT D816V, FLT3 D835Y, and PDGFRA D842V are activation loop mutations commonly found in acute leukemias, gastrointestinal stromal tumors, and other cancer types. They are highly oncogenic, but their mechanisms of activation are not well understood (27, 29-31).

Given the therapeutic importance of kinases in oncology, our deep understanding of kinase structure-function relationships, and the need for consistent data across mutations for NGS analysis, we generated a structural alignment of all missense mutations in the kinase and JM domains of 48 RTKS based on mutations reported in the Catalogue of Somatic Mutations in Cancer (COSMIC), the largest curated database of clinical cancer mutations (Figure 1C) (32). The results were functionally annotated and entered into a database that can be used as a tool to evaluate novel RTK missense mutations based on their structural position relative to known activating missense mutations. The database can be accessed either in full through GitHub (https://github.com/quantumdolphin/kinase_paper) or via a user-friendly web application (https://kinase-mutation-atlas.streamlit.app). We provide experimental data validating the use of this tool by showing several VUSs that structurally align with oncogenic mutations in other kinases are activating. Specifically, we demonstrate in a cell-based model that several VUSs in FLT3 and PDGFRA lead to kinase activation. Lastly, we show how this database can be used to develop testable mechanistic hypotheses regarding previously uncharacterized mutations in several kinase regions.

## Methods

### Data collection

Forty eight RTKs mutated in cancer were selected (Table S1) (32). A multiple sequence alignment of kinase and JM domain sequences was generated using MUSCLE (Appendix 1) (33). Mutational data and corresponding protein sequences were downloaded from the Catalogue of Somatic Mutations in Cancer (COSMIC) (32). There were 5778 residue positions with mutations (Figure S2). Mutations at 300 positions had annotations in OncoKB, a precision oncology knowledge database that classifies mutations as gain-of-function (GOF), likely gain-of-function (LGOF), loss-of-function (LOF), likely loss-of-function (LLOF), neutral, and likely neutral (34, 35). These annotations were added to the database.

Thirty-six of the 48 kinases had kinase domain structures in the Protein Data Bank with <2.7 Å resolution, which we selected as a cut-off for adequate resolution (36, 37). Many kinases had multiple structures therefore for each kinase a representative structure was selected that had: (1) maximum coverage of kinase and JM domains, and (2) high quality (high resolution <2.7 Å and low R-free, a crystallographic quality metric). For the 12 kinases without structures, homology models were obtained from the Kinametrix web server (38-40).

UniProt numbering obtained from the PDBrenum server was used for representative structures (41, 42). In cases with multiple transcripts in Uniprot and/or COSMIC, transcripts for the crystal structure were used.

### Structure-based clustering

Custom Python scripts and the MDAnalysis package were used for clustering and analysis (code and data: https://github.com/quantumdolphin/kinase_paper) (43, 44). Mutations from COSMIC were mapped to the three-dimensional structure using side chain center of masses for subsequent clustering. Hierarchical clustering was conducted using the Euclidean distance and Ward linkage method, with the X, Y, and Z coordinates of the side chain center of masses as input parameters. The top 10 clusters that maximized GOF mutations and minimized LOF mutations were selected for further analysis (Figures S3). For further details on clustering see Supplemental Methods. ChimeraX was used for visualization (45). A user-friendly, publicly available web application was built using Streamlit that allows the user to input a kinase mutation and output a list of mutations in other kinases that are either structurally aligned or in close proximity to the initial mutation (https://kinase-mutation-atlas.streamlit.app) (46).

### Ba/F3 Cell Assays

Ba/F3 cells were maintained in RPMI 1640 (Cellgro) supplemented with 1% penicillin-streptomycin (PS), murine IL-3 and 10% fetal bovine serum (FBS). pHAGE-PDGFRA (Addgene) and pLVX-Puro-FLT3-eBFP were used for generating described mutations using a site directed mutagenesis kit (Agilent) (17, 20). For primers, see Supplemental Methods. HEK293T cells were transfected with 8 μg of either PDGFRA or FLT3 vector, 2 μg of pCMV-VSVG, and 4 μg of PAX2 vectors using the X-tremeGENE HP DNA transfection agent (Roche #6366244001). The viral supernatants were collected after 48 hours and used for Ba/F3 cell transduction. Three million cells were infected with lentiviral supernatant and polybrene (Santa Cruz Biotechnology #SC-134220) using spinfection followed by incubation for 5 hours at 37°C. After infection, cells were selected using 1μg/ml puromycin (Invitrogen #ant-pr-1). Following selection, IL-3 withdrawal was performed, and viability assessed in IL-3 free media.

## Results

### Top mutation clusters are within key regulatory regions

The top 10 clusters of mutations that maximized GOF and minimized LOF mutations were concentrated in key regulatory regions close to the active site (Figure 1C). These regulatory regions include the activation loop, the glycine-rich loop (G-loop), the JM domain, the αC-helix, and the loops N-terminal and C-terminal to the *α*C-helix (the *β*3-*α*C loop and the *α*C−*β*4 loop). To demonstrate the utility of our database as a tool, we describe the analysis of a sample of mutations from several of the top clusters. The results are organized based on the regulatory regions in which the sample mutations are found.

### The activation loop is a key site of activating mutations

The activation loop recognizes and binds substrate, undergoing large conformational changes between inactive and active states, with phosphorylation stabilizing the active state. Several well-characterized GOF mutations were found within the activation loop (Figure 2A,B), consistent with known literature (27, 47, 48). Mutations were absent from the N-terminal portion of the activation loop, called the DFG motif (Figure 2C), likely because the DFG is critical for catalysis and DFG mutations would be predicted to be LOF. However, many mutations, including VUSs, were found in the ∼10 amino acids immediately following the DFG, and, to a lesser extent, portions of the protein surrounding/contacting this area of the activation loop. Two aspartic acid (Asp) residues in the activation loop (positions 7 and 11, Figure 2B-D) had very high numbers of GOF mutations in the type 3 RTKs: FLT3, KIT, and PDGFRA (Figure 2D). These Asp residues sit on either end of a small helix known as a 3-10 helix, which we hypothesize may be involved in the mechanism of activation by the Asp mutations. The Asp at position 7 is the Asp^*β*9^ (FLT3 D835, KIT D816, and PDGFRA D842) which is recurrently mutated in many cancer types (27). Only the type 3 RTKs had large numbers of mutations that structurally aligned to this site (Figure 2D), however there were small numbers of mutations at this position in several other kinases, including KDR D1052X mutations that have been shown to increase KDR activity in a tissue culture model (49). Though these KDR mutations are not annotated as GOF in OncoKB, we hypothesize that based on their structural alignment with other highly oncogenic, recurrent mutations in KIT, PDGFRA, and FLT3, that they are in fact GOF.

**Figure 2.**
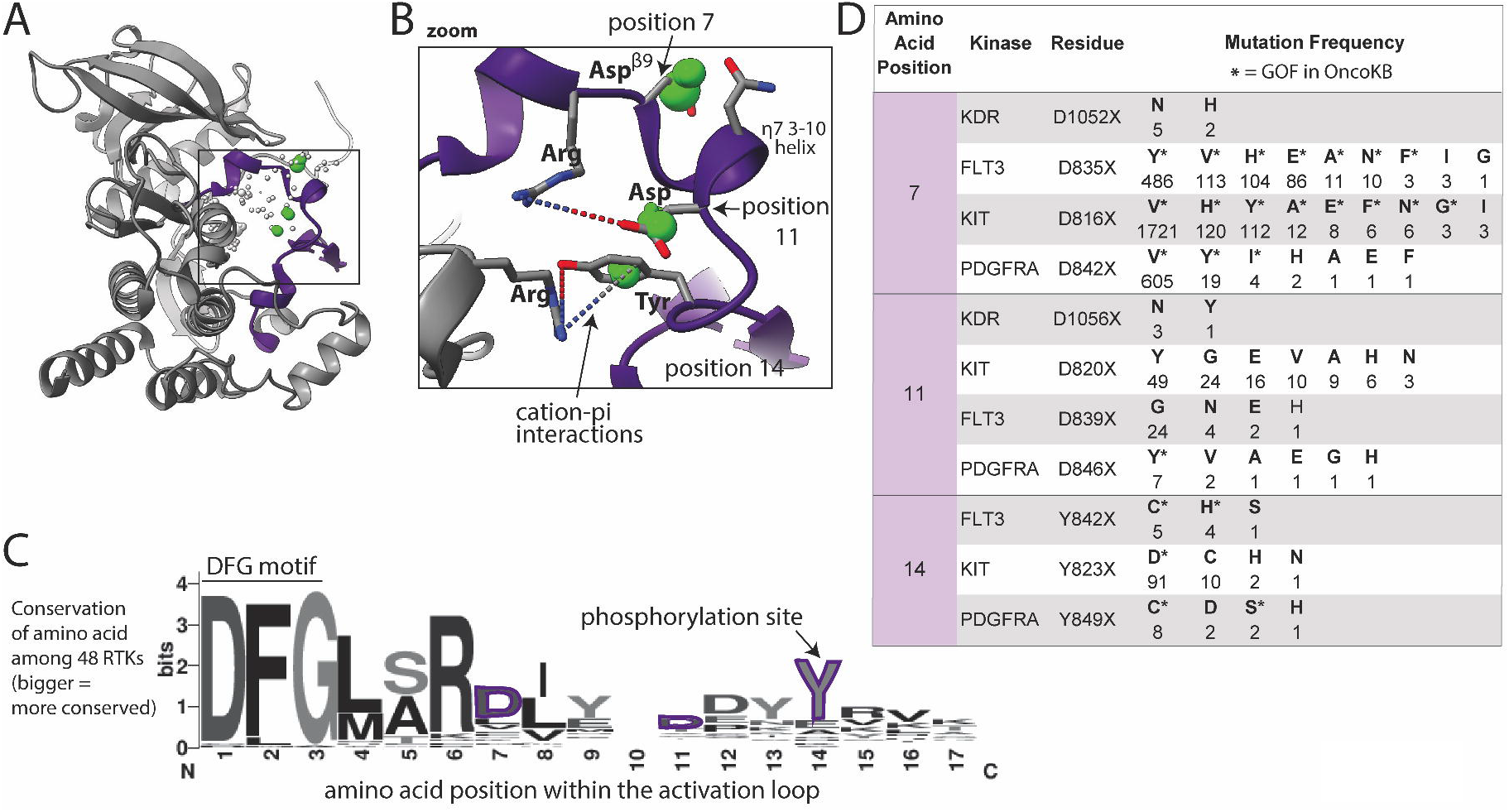
Activation loop mutations are common. A) Activation loop (purple) shown in KIT, as a representative structure; a subset of residues with reported mutations are shown as dots, those highlighted in the text are green, the size of the dots indicate number of mutations at that site across the 48 kinases as reported in COSMIC, with numbers shown in panel D. B) Zoom in of areas of frequent mutation in the N-terminal portion of the activation loop. Side chains are shown for discussed residues (gray = carbon, blue = nitrogen, red = oxygen). C) Sequence logo based on alignment of the N-terminal portion of the activation loop in all 48 kinases, starting with the DFG motif; the larger the letter, the more conserved the amino acid at that position across the 48 kinases (bits on the y-axis is a measure of the degree of conservation). D) Frequency and identity of a sample of mutations at highlighted positions in the N-terminal portion of the activation loop. The amino acid position correlates with the positions labeled in panels B and C. The starred mutations are GOF in OncoKB.

The Asp at the C-terminal end of the 3-10 helix in KIT, KDR, PDGFRA, and FLT3 (Figure 2B, position 11) is also frequently mutated, but less well characterized than Asp^*β*9^. We hypothesize that most mutations at this position will be activating by disrupting key electrostatic interactions (dotted lines between R815 and D820 in the case of KIT shown in Figure 2B) that hold the activation loop in the inactive conformation. Additionally, there is a tyrosine (Tyr) (position 14) phosphorylation site C-terminal on the activation loop that is highly mutated. Given that this Tyr is a phosphorylation site, mutations may be phospho-mimetics as several of the mutant residues are Asp, which has a negative charge resembling a phosphate group.

### Mutations in the G-loop may impact function by changing loop flexibility

A large number of mutations were located around the G-loop, which is characterized by the consensus sequence (GXGXΦG), where Φ is usually a Tyr or phenylalanine (Phe) (Figure 3A-D) (22, 50). The G-loop is highly flexible, positioning ATP for phosphoryl transfer in the active site.

**Figure 3.**
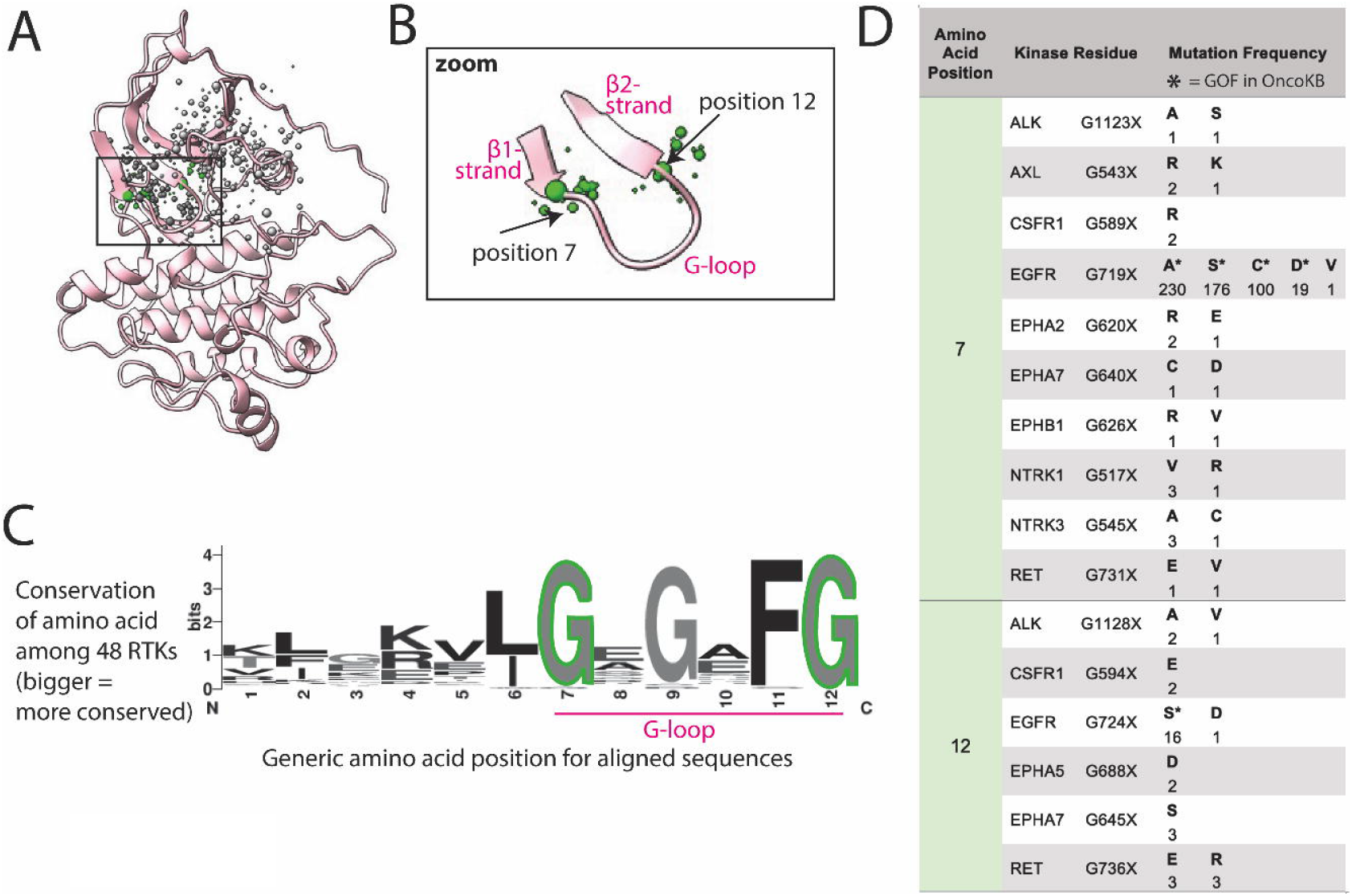
G-loop mutations are at the hinges of the loop. A) G-loop shown in FLT3, as a representative structure; a subset of residues with reported mutations are shown as dots, the residues highlighted in the text are green, the size of the dots indicate number of mutations at that position. B) Zoom of areas of frequent mutation in the G-loop. C) Sequence logo based on alignment of G-loop in all 48 kinases; the larger the letter, the more conserved the amino acid at that position. D) Frequency and identity of a sample of mutations that occur at each highlighted position in the G-loop, the amino acid position correlates with the positions labeled in panels B and C. The starred mutations are annotated as GOF in OncoKB.

Mutations reported in COSMIC within the G-loop are almost exclusively at the first and last glycine (Gly), positions 7 and 12, at the two ends of the loop (Figure 3B). Though mutations in EGFR at these positions are well characterized as GOF, there are uncharacterized mutations in at least 10 other RTKs at these positions (Figure 3D) (51). Gly, lacking a side chain, plays a unique role in protein flexibility, and the first and last Gly in the G-loop act as hinge points for the movement of the loop. Mutating these to any other amino acid is predicted to decrease loop flexibility, potentially locking it into an active-like conformation. Thus, we hypothesize that all mutations of these 2 critical glycines could impact loop flexibility and thus activation, as previously demonstrated for G-loop mutations in EGFR (52).

Within the same cluster, there are also numerous mutations in the *β*_1_ and *β*_2_ strands of the N-lobe flanking the G-loop, which could also plausibly, but more speculatively, be activating through modifying the structure and/or dynamics of the G-loop.

### Mutations in the JM domain likely disrupt autoinhibitory interactions with the kinase domain

The *α*C-helix, which abuts the JM domain in some kinases, is important in kinase regulation and undergoes significant conformational change between active and inactive states (Figure 1) (22). The *α*C-helix contains a conserved glutamate (Glu) that faces inward and helps align the catalytic lysine (Lys) necessary for phosphoryl transfer (Figure S1, Figure 4). This conserved Glu, and other amino acids on the inward-facing side of the *α*C-helix, have few mutations. However, mutations of outward-facing amino acids on the αC-helix that interact with the JM domain are more common (Figure 4A-D).

**Figure 4.**
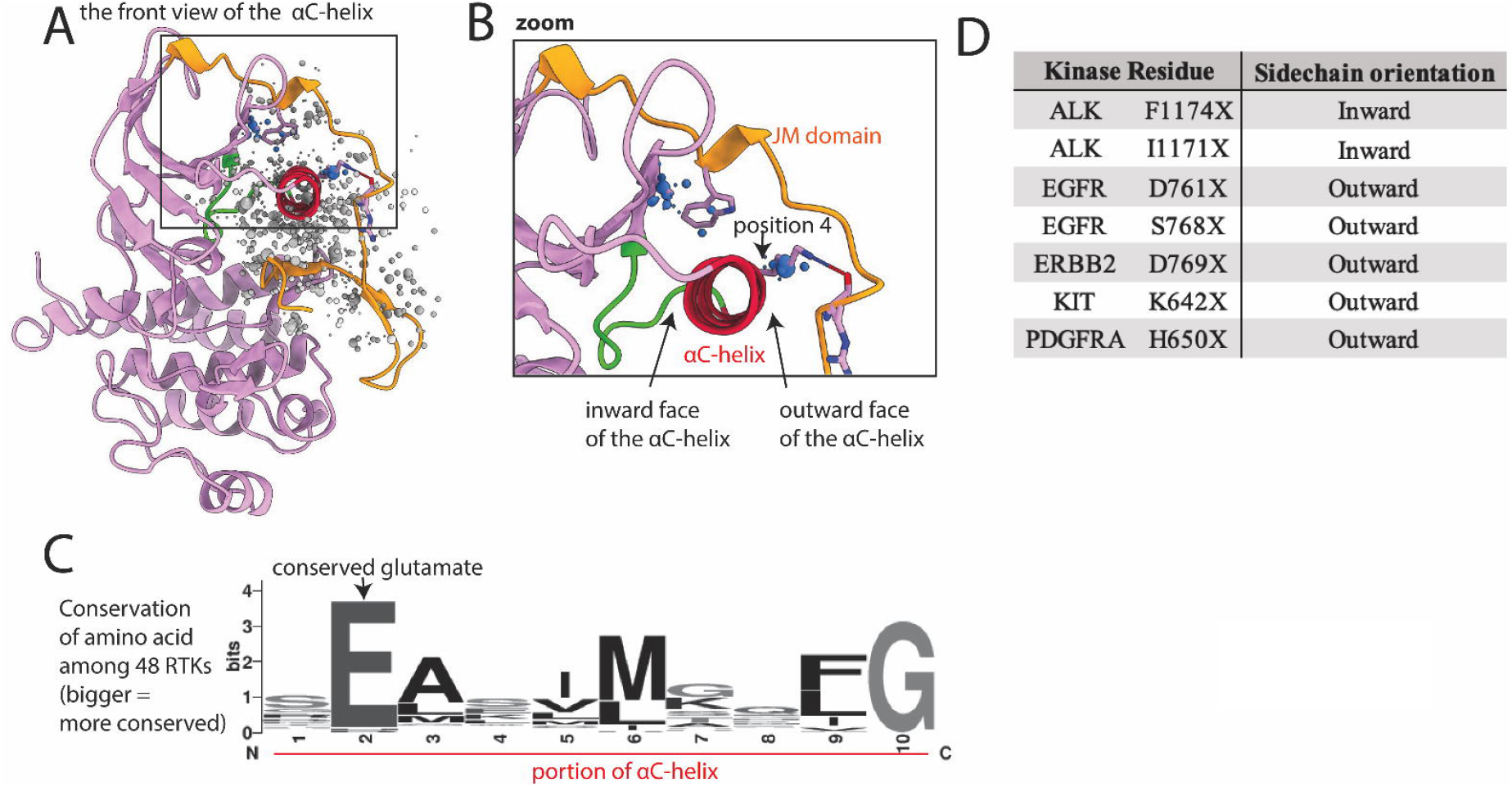
Mutations cluster at the interface between the kinase domain and juxtamembrane (JM) domain. A) Kinase domain/JM domain interface shown in EGFR, as a representative structure; the kinase domain is pink, with the αC-helix highlighted in red and the *α*C−*β*4 loop highlighted in green, and the JM domain is gold; a subset of residues with reported mutations are shown as dots, the residues highlighted in the text are blue. B) Zoom of highlighted areas in the kinase-JM domain interface; position 4 is a residue discussed in the text that we hypothesize is vulnerable to mutational activation based on its position between the αC-helix and the JM domain. C) Sequence logo of a portion of the αC-helix based on alignment of all 48 kinases; the larger the letter, the more conserved the amino acid at that position. D) Oncogenic mutations on the inward face of the αC-helix are uncommon compared to those on the outward face of the αC-helix, which interacts with the JM domain.

The JM domain has autoinhibitory interactions with the kinase domain in the inactive state, especially in type 3 RTKs. There are numerous well-characterized JM domain GOF mutations that abolish JM-to-kinase domain autoinhibitory interactions. We found 2 large areas of mutations involving the JM domain and the contacting kinase domain, including outward-facing portions of the αC-helix (Figure 4A,B). KIT K642E is an example of a characterized GOF mutation previously identified on the outward face of the αC-helix (53). K642 in wildtype KIT forms a hydrogen bond with the backbone of the JM domain; therefore mutating it would break this interaction. We hypothesize that mutations in the analogous Lys of other kinases, such as FLT3 (shown in Figure 4B, position 4) and PDGFRA, will also be activating. Many other mutations in the interface between the JM and kinase domains may also be GOF by disrupting autoinhibition. For example, we previously reported that a kinase domain VUS, FLT3 A680V, which sits at this interface was activating (17). As this mutation is on the interface between JM and kinase domains, the mechanism of activation may involve disruption of autoinhibition.

### Mutations in the *α*C−*β*4 loop impact kinase function through allostery

In addition to mutations on portions of the *α*C-helix interacting with the JM domain, there is also a large number of mutations in the 2 loops flanking the αC-helix. These may lead to GOF by changing orientation of the helix. On one side of the αC-helix is the *β*3-*α*C loop, which harbors well characterized mutations in BRAF, EGFR and HER2 (28). On the other side of the helix is the *α*C−*β*4 loop (Figure 5A-C) which is involved in long-range allosteric coupling with the catalytic kinase core by orchestrating movement of the *α*C-helix with kinase activation (54). This *α*C−*β*4 loop has a large number of mutations in COSMIC, though relatively few have been characterized as activating (Figure 5D).

**Figure 5.**
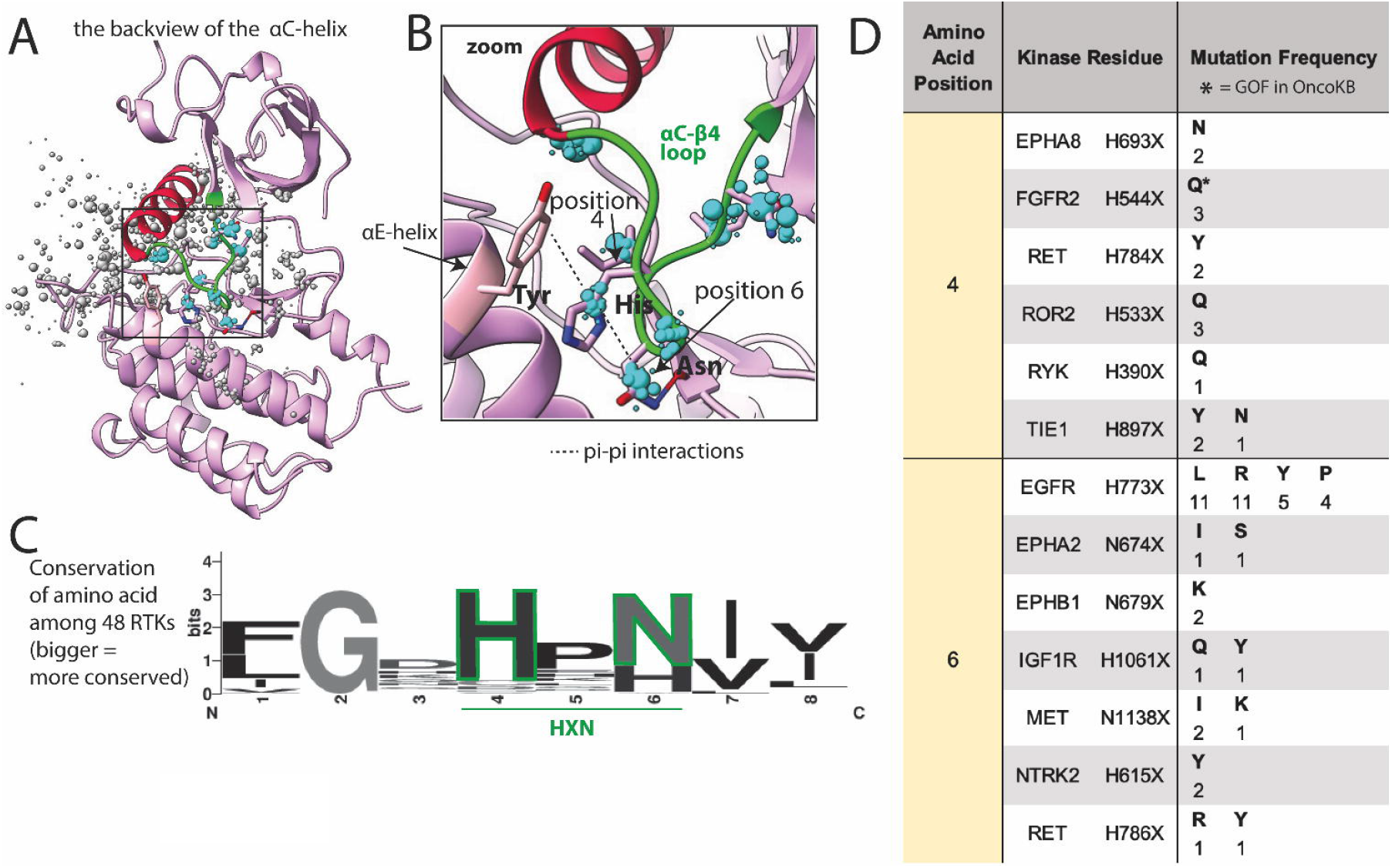
Common mutations in the *α*C−*β*4 loop. A) *α*C−*β*4 loop (green) shown in FGFR2, as a representative structure; a subset of residues with reported mutations are shown as dots, the residues highlighted in the text are blue, the size of the dots indicate number of mutations at that position. B) Zoom of areas of frequent mutation in the *α*C−*β*4 loop. Side chains are shown for residues discussed in the text (pink = carbon, blue = nitrogen, red = oxygen). C) Sequence logo based on alignment of the *α*C−*β*4 loop in all 48 kinases; the larger the letter, the more conserved the amino acid at that position. D) Frequency and identity of a sample of mutations that occur at each highlighted position in the *α*C−*β*4 loop, the amino acid position correlates with the positions labeled in panels B and C. The starred mutations are annotated as GOF in OncoKB.

### Uncharacterized mutations that align with known oncogenic mutations are activating

We hypothesized that our database of structurally aligned mutations could be used to help identify previously uncharacterized mutations as activating. A set of 5 PDGFRA mutations and 5 structurally analogous FLT3 mutations, including 2 activation loop mutations, 2 *α*C−*β*4 loop mutations, and 6 JM domain mutations that interact with the kinase domain at different locations were selected for experimental validation (Figure 6A,B). These mutants were expressed in Ba/F3 cells, which become cytokine-independent in the presence of an activated kinase and have been used extensively to evaluate for kinase activation (55). All selected mutations were within key regulatory regions, though only 3 (PDGFRA Y849C, V561D, Y555C) were designated GOF in OncoKB (Figure 6B). PDGFRA D842V and FLT3 D835Y mutations were used as positive controls.

**Figure 6.**
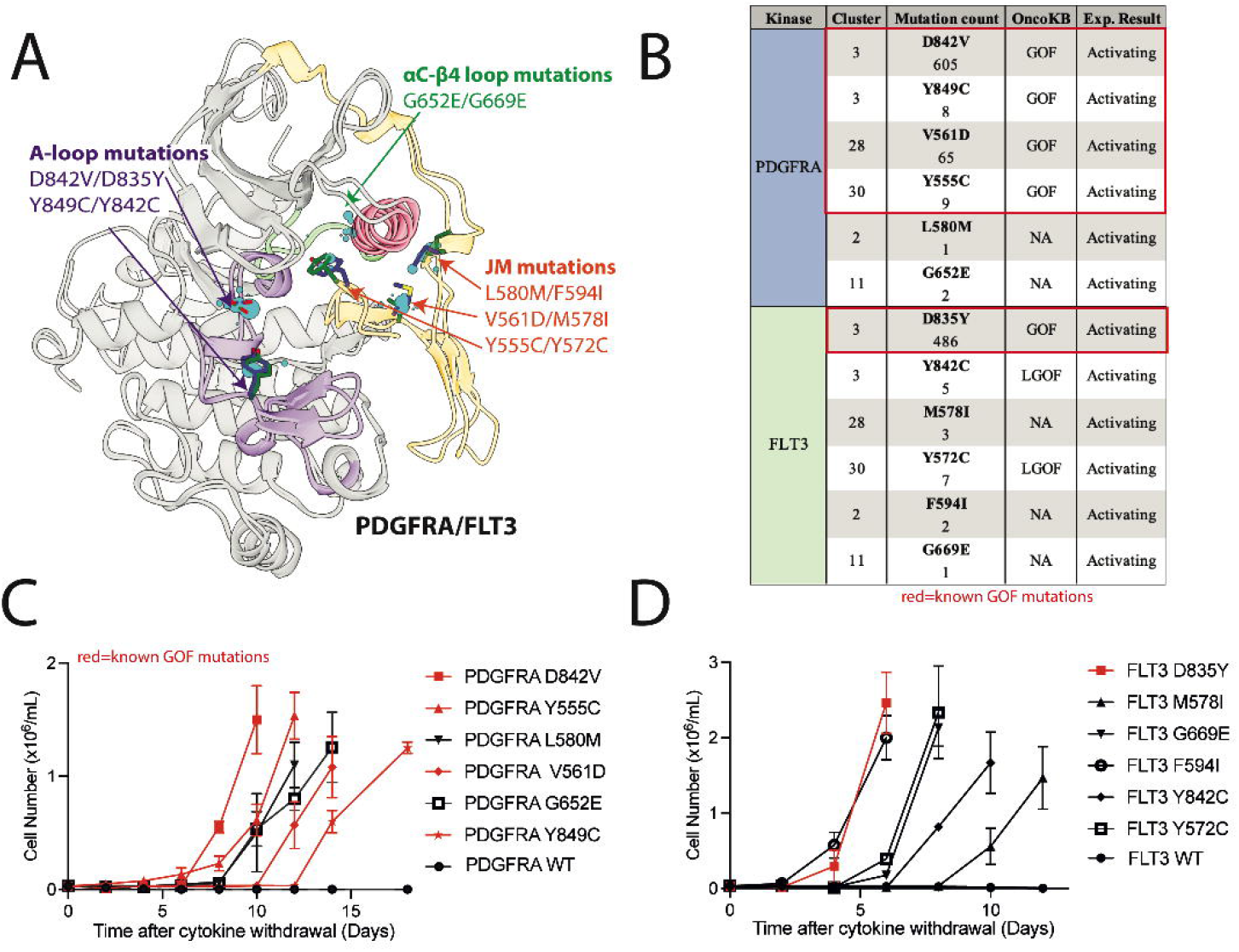
Previously uncharacterized FLT3 and PDGFRA mutations in areas of high mutational burden are activating. A) Alignment of kinase and JM domain structures of PDGFRA and FLT3 in the inactive conformation with evaluated mutations highlighted. B) List of mutations with OncoKB annotations. IL-3 independent growth of Ba/F3 cells expressing C) PDGFRA and D) FLT3 mutations.

Two of the selected FLT3 mutations were designated LGOF in OncoKB but had been previously shown to be activating in tissue culture models. These mutations include, 1) the activation loop mutation FLT3 Y842C, which is structurally analogous to the PDGFRA GOF mutation Y849C, and, 2) FLT3 Y572C, which is structurally analogous to the PDGFRA GOF mutation Y555C (56-58). All of these mutations were also activating in our Ba/F3 assays (Figure 6C,D).

Two of the selected PDGFRA mutations (L580M and G652E) and 3 of the selected FLT3 mutations (M578I, F594I, and G669E) had no annotations in OncoKB and were not characterized in the literature. However, these mutations were in top clusters and within regulatory regions of high mutational burden. When expressed in Ba/F3 cells, we found all 5 mutations were activating (Figure 6C,D). These data support the use of this database as a resource in the analysis of kinase mutations found in clinical NGS.

## Discussion

With the explosion of next-generation sequencing data and the concurrent availability of protein structures through both the Protein Data Bank and models generated by programs such as AlphaFold, we have an unprecedented opportunity to use protein structure to develop tools to understand the biological impact of clinical cancer mutations (18, 36). Here, we describe the generation of a database of structurally aligned and functionally annotated cancer mutations across 48 kinases that can be used as a resource to analyze kinase VUSs, with potential clinical implications to guide targeted therapy. We have focused our analysis on clusters of mutations centered on 4 regions of the kinase domain that are known to be important for regulating activity, and thus enriched in both known GOF mutations and numerous VUSs that we propose are also likely to be activating:

1. **Activation loop**. Here, we found mutations absent from the N-terminal portion of the activation loop, called the DFG motif. In the Ser/Thr kinase, BRAF, oncogenic, inactivating DFG mutations are well described (59, 60). These DFG-mutant BRAF proteins activate cell signaling by causing the inactive BRAF to heterodimerize and transactivate CRAF (59, 60). However, inactivating DFG mutations that activate oncogenic signaling have not been described in other kinases. The lack of DFG mutations in our database was expected given that mutation of the DFG prevents kinase catalytic function. In contrast, mutations of Asp^*β*9^, which occupies a central position on the activation loop and is highly mutated in cancer (e.g. KIT D816V, PDGFRA D842V, FLT3 D835Y, position 7 in Figure 2), comprised one of the top mutation clusters, providing proof of principle that structural areas of high mutational burden across kinases harbor oncogenic mutations. Asp^*β*9^ mutations that introduce hydrophobic or aromatic side chains (Val, Tyr, Ala, Phe, Ile) are known to be activating (20, 61, 62). Even a conservative Asp to Glu substitution, where the Glu differs from the wildtype Asp by only a single methyl group, can be activating (e.g. PDGFRA D842E, FLT3 D835E) indicating precise charge placement is required to stabilize the inactive state (20, 62). Though the mechanism by which Asp^*β*9^ mutations activate kinases is not well understood, our analysis of the structures of multiple kinases simultaneously suggests the orientation and position of the Asp^*β*9^ may stabilize the charge distribution (or the dipole moment) of the 3-10 helix (Figure 2B), which could preserve the inactive conformation. Therefore, mutation of the Asp^*β*9^ could shift the charge distribution and destabilize the inactive state. Several other mutations in the activation loop C-terminal to the Asp^*β*9^ likely have a similar mechanism, i.e., destabilizing the inactive state, such as the Y842C mutation in FLT3, which we confirm to be activating in Ba/F3 cells.
2. **G-loop**. Mutations in this key structural and functional element have been previously described in EGFR in non-small cell lung cancer, in BCR::ABL1 in chronic myelogenous leukemia, and in BRAF in colon cancer (51, 63, 64). Interestingly, the EGFR and BCR::ABL1 mutations confer poor prognosis with aggressive, drug-resistant disease (51, 63). Other G-loop mutations have been reported, but with unknown prognostic significance. For example, ALK G1128A, a G-loop mutation, has intermediate oncogenic activity compared to other ALK mutations, but there is not enough data to determine prognostic significance (21). Our analysis showed mutations within the G-loop, including the EGFR and ALK mutations discussed above, are located almost exclusively at the first and last Gly residues at the two ends of the loop. Analogous mutations have been reported in at least 10 other RTKs, including RET, NTRK1/3, and several ephrin receptors, albeit with low prevalence (Figure 3D). We hypothesize that these increase kinase activity by rigidifying the loop at the critical glycine hinge-points, potentially locking the loop into an active-like conformation. There are also numerous mutations in the short *β*_1_ and *β*_2_ strands that flank the G-loop. It is possible that some of these could also be activating through modulating the structure and/or dynamics of the G-loop, but little information is currently available to support this hypothesis.
3. **JM domain**. Interactions between the JM and kinase domains stabilize the autoinhibited states, and thus mutations in amino acids involved in these interactions are frequently activating. Such mutations include well-characterized GOF mutations on the JM domain itself, such as Y555C in PDGFRA, and mutations on the ‘outward-facing’ portion of the αC-helix that interact with the JM domain, such as K642E in KIT. We identified additional clusters of uncharacterized mutations at the interface between the JM and αC-helix that we postulated would also destabilize the autoinhibited state and confirmed that L580M in PDGFRA and M578I and F549I in FLT3 were in fact activating in Ba/F3 cells.
4. The 2 loops flanking the αC-helix, the ***β*3-*α*C loop** and the ***α*C−*β*4 loop**. It has been shown that small deletions in the *β*3-*α*C loop change the orientation of the αC-helix, leading to activation (28). We hypothesize that missense mutations in the *α*C−*β*4 loop also alter the orientation of the αC-helix and can cause kinase activation. This hypothesis is based on literature showing the *α*C−*β*4 loop (Figure 5A-C) is involved in long-range allosteric coupling with the catalytic kinase core and can orchestrate movement of the *α*C-helix with kinase activation (54). In particular, the HXN motif (X=any amino acid) is the region of the *α*C−*β*4 loop that forms the pivot point for *α*C-helix movement (Figure 5B). GOF mutations have been identified in the HXN as well as in other portions of the *α*C−*β*4 loop. Substitution of the histidine (His) within the HXN, such as the GOF mutation FGFR2 H544Q (Figure 5B, C, position 4), eliminates a favorable pi-pi stacking interaction between the His and a conserved Tyr in the αE-helix (Figure 5B), potentially altering the structure of the inactive state (65). In addition to FGFR2, mutations involving the His have been reported in EPHA8, RET, ROR2, RYK, and TIE1 (Figure 5D), and we predict these are similarly activating by eliminating pi-pi stacking interactions and changing the conformation of the *α*C−*β*4 loop and the αC-helix. The Asn (Figure 5B, C, position 6) of the HXN motif is less well conserved but the Asn side chain is hypothesized to be part of a water-mediated hydrogen bond network aiding allosteric communication between the *α*C-helix and the catalytic core (66). Substitution of Asn with amino acids that change the hydrogen bonding network can affect allosteric signaling and impact kinase activation. Consistent with this assertion, EGFR has a known GOF mutation (EGFR H773L) in this position that eliminates hydrogen-bonding (67). IGF1R, NTRK2, RET, EPHA2, EPHB1, and MET also have mutations in this position, which we predict to be activating (Figure 5D). Overall, the *α*C−*β*4 loop appears to be highly susceptible to GOF mutations, and most mutations in this loop, even seemingly conservative ones, seem likely to be GOF. To begin to explore this hypothesis, we tested VUSs in PDGFRA and FLT3 at another position in the loop, 2 residues N-terminal to the HXN motif, and both (G652E and G669E, respectively) were activating in Ba/F3 cells.

In summary, there are a large number of kinase VUSs that align structurally with characterized kinase mutations. Given the complexity of trying to predict protein function based on point mutations, oncologists and tumor boards are likely to continue relying on integrated information from multiple sources to interpret clinical NGS and make molecularly targeted therapy recommendations. Here, we provide a database of structurally aligned and functionally annotated mutations across 48 RTKs that can be used as a consistent resource across RTKs to evaluate VUSs for structural alignment with known activating mutations, while also providing insight into potential mechanisms of activation and regulation. Overall, we expect our database to be an important addition to the current tools and resources used to analyze clinical NGS.

## Acknowledgements

This work is generously supported by Julia’s Legacy of Hope St. Baldrick’s Foundation Consortium Research Grant (YP) and the Bachrach Family Foundation (BAW).

## Data availability statement

The code and data generated in this study are publicly available: https://github.com/quantumdolphin/kinase_paper

