## Supplemental Material for "A structure-based tool to interpret the significance of kinase mutations in clinical next generation sequencing in cancer"

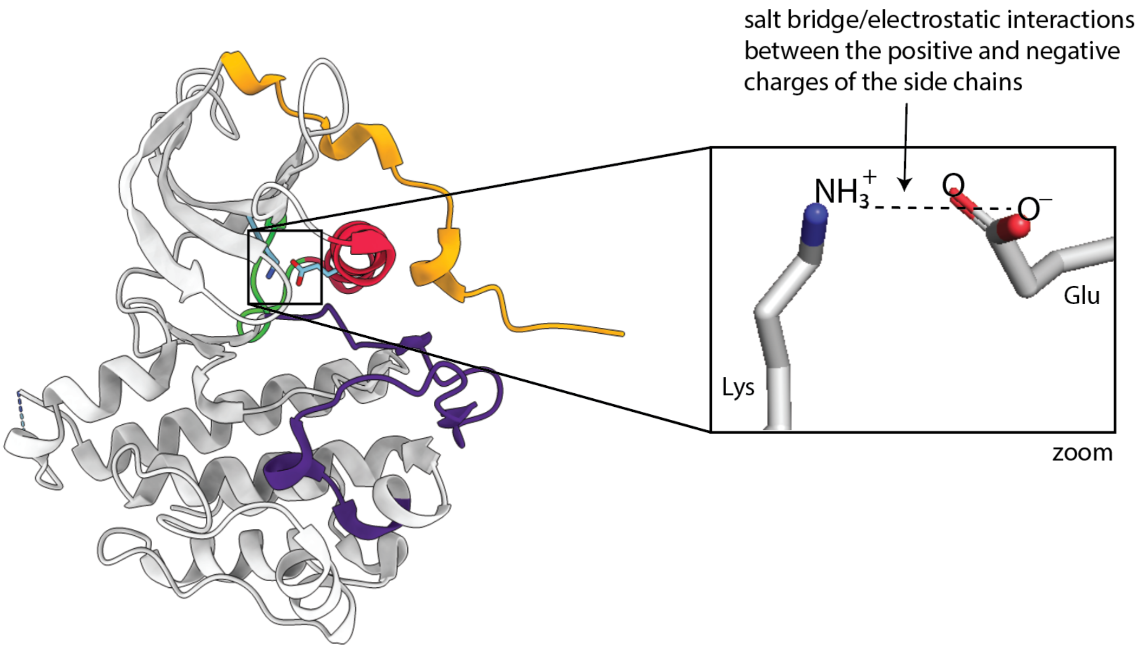


**Figure S1. Electrostatic interactions between the αC-helix and the kinase core.** This figure shows the salt bridge that is formed between the Glu on the αC-helix and the Lys in the core of the kinase. This salt bridge is essential for the catalytic activity of the kinase and enables phosphoryl transfer.

**Table S1: Representative structures and the corresponding COSMIC data**

| Gene_name | Representative structure | Uniprot_id | RTK_Family | COSMIC_ID |
|---|---|---|---|---|
| ALK | 2XP2 | Q9UM73 | ALK | COSG350 |
| AXL | homology model | P30530 | AXL | COSG10854 |
| CSF1R | 3KRJ | P07333 | PDGFR | COSG107 |
| DDR1 | 6FTO | Q08345 | DDR | COSG9377 |
| DDR2 | homology model | Q16832 | DDR | COSG139 |
| EGFR | 1M14 | P00533 | EGFR | COSG150 |
| EPHA1 | homology model | P21709 | EPH | COSG152 |
| EPHA10 | homology model | Q5JZY3 | EPH | COSG37048 |
| EPHA2 | 5EK7 | P29371 | EPH | COSG155 |
| EPHA3 | 2QOQ | P29320 | EPH | COSG154 |
| EPHA4 | homology model | P54764 | EPH | COSG153 |
| EPHA5 | 2r2p | P54756 | EPH | COSG22150 |
| EPHA7 | 2REI | Q15375 | EPH | COSG158 |
| EPHA8 | homology model | P29322 | EPH | COSG159 |
| EPHB1 | 5MIA | P54762 | EPH | COSG160 |
| EPHB2 | 3ZF7 | P29323 | EPH | COSG40191 |
| EPHB3 | 5L6P | P54753 | EPH | COSG162 |
| EPHB4 | 6FNK | P54760 | EPH | COSG163 |
| EPHB6 | homology model | O15197 | EPH | COSG19012 |
| ERBB2 | 3PP0 | P04626 | EGFR | COSG165 |
| ERBB3 | homology model | P21860 | EGFR | COSG166 |
| ERBB4 | 3BCE | Q15303 | EGFR | COSG167 |
| FGFR1 | 3AM6 | P11362 | FGFR | COSG171 |
| FGFR2 | 3B2T | P21802 | FGFR | COSG173 |
| FGFR3 | 6PNX | P22607 | FGFR | COSG97353 |
| FGFR4 | 4QF5 | P22455 | FGFR | COSG1767 |
| FLT1 | 3HNG | P17948 | VEGFR | COSG187 |

| | | | | |
|---|---|---|---|---|
| FLT3 | 1T3 | P36888 | PDGFR | COSG210 |
| IGF1R | 5FXQ | P08069 | InsR | COSG215 |
| INSR | 1IRK | P06213 | InsR | COSG220 |
| IRR | homology model | P14616 | InsR | COSG219 |
| KDR | 4ASD | P35968 | VEGFR | COSG615273 |
| KIT | 5GOE | P10721 | PDGFR | COSG211 |
| LTK | homology model | P29376 | ALK | COSG91303 |
| MERTK | 7AB2 | Q12866 | Axl | COSG106715 |
| MET | 2G15 | P08581 | Met | COSG12 |
| MST1R | 3PLS | Q04917 | Met | COSG315 |
| NTRK1 | 6PL4 | P04629 | Trk | COSG336 |
| NTRK2 | 4AT5 | Q16620 | Trk | COSG27120 |
| NTRK3 | 4YMJ | Q16288 | Trk | COSG9247 |
| PDGFRA | 5K5X | P16234 | PDGFR | COSG13 |
| RET | 6NEC | P07949 | Ret | COSG17 |
| ROR1 | 6T9V | Q01973 | Ror | COSG16 |
| ROR2 | 3ZZW | Q01974 | Ror | COSG417 |
| RYK | 6TUA | P34925 | Ryk | COSG620600 |
| TEK | 1FVR | Q02763 | Tie | COSG488 |
| TIE1 | homology model | P35590 | Tie | COSG494 |
| TYRO3 | homology model | Q06418 | Axl | COSG515 |

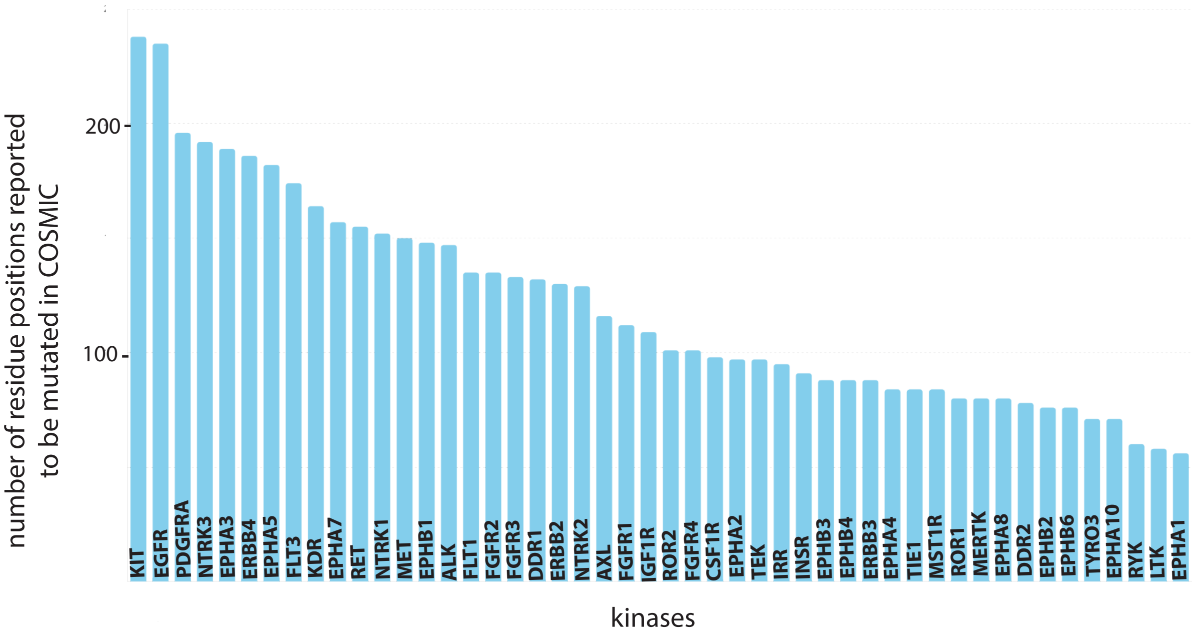


**Figure S2.** The distribution of residue positions with mutations across receptor tyrosine kinases. The dataset has a total of 5778 residue positions from 48 receptor tyrosine kinases.

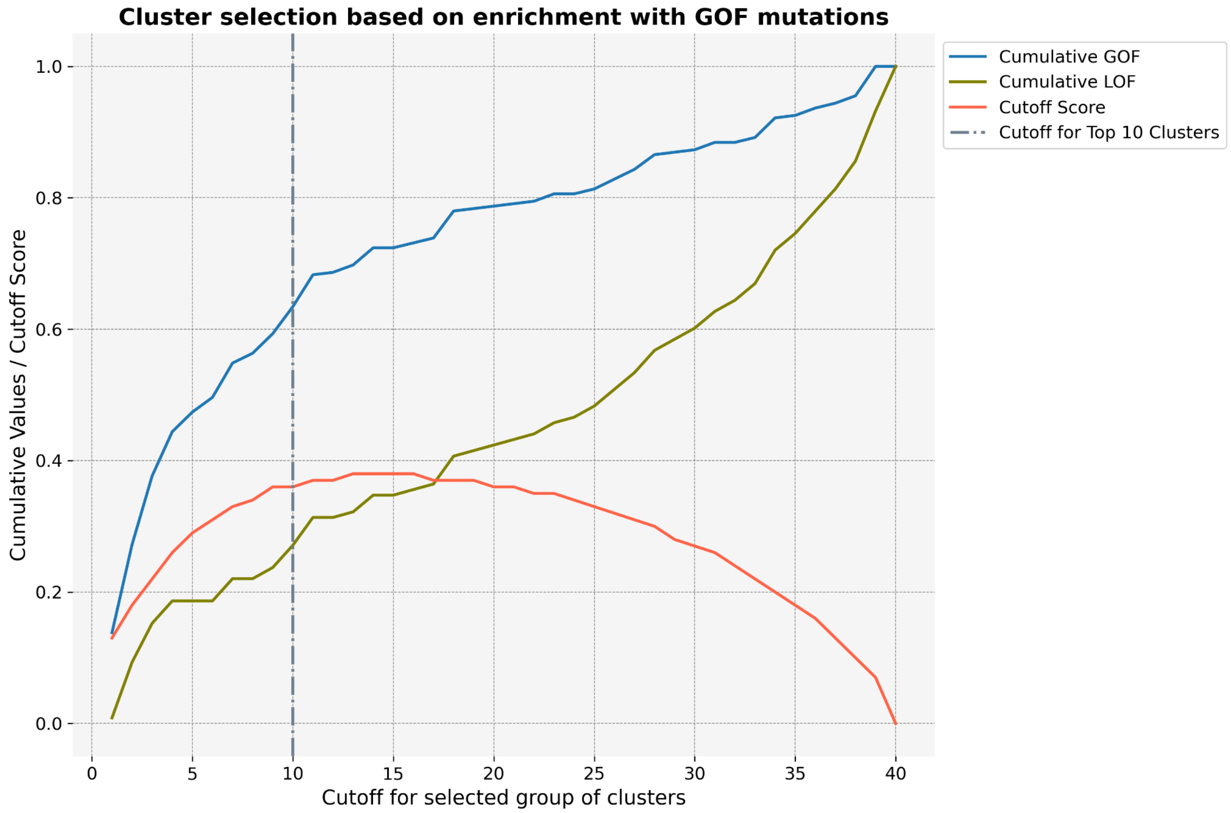


**Figure S3.** Ten clusters that maximized GOF mutations and had few LOF mutations were selected as the top ten clusters.

### Supplemental Methods

#### Clustering

Hierarchical clustering was conducted using the Euclidean distance and Ward linkage method, with the X, Y, and Z coordinates of the side chain center of masses as the input parameters. We chose 40 clusters empirically. When generating fewer clusters, e.g., 20, the clusters encompass multiple key functional regions such as the activation loop and αC-helix. Using numbers of clusters larger than 40 generated very small clusters with only a few mutations, making it difficult to perform further analyses.

Next, a subset of clusters that were enriched in gain-of-function (GOF) mutations and minimized loss-of-function (LOF) mutations, were selected. The 'net_GOF' and 'net LOF' mutations for each cluster were calculated based on OncoKB annotations as the sum of GOF and likely GOF annotations, or LOF and likely LOF values, respectively as shown in eq 1,2.

$$\text{net GOF}_{(i)} = \text{GOF}_{(i)} + \text{likely GOF}_{(i)} \quad (1)$$

$$\text{net LOF}_{(i)} = \text{LOF}_{(i)} + \text{likely LOF}_{(i)} \quad (2)$$

These values were normalized to the total GOF or LOF ('norm GOF' and 'norm LOF'), as follows:

$$\text{norm GOF}_{(i)} = \frac{\text{net GOF}_{(i)}}{\text{total GOF}} \quad (3)$$

$$\text{norm LOF}_{(i)} = \frac{\text{net LOF}_{(i)}}{\text{total LOF}} \quad (4)$$

where,

$$\text{total GOF} = \int_{i=1}^{40} \text{net GOF}_{(i)} \quad (5)$$

$$\text{total LOF} = \int_{i=1}^{40} \text{net LOF}_{(i)} \quad (6)$$

Cumulative values for GOF and LOF were calculated as shown in equations 7 and 8. A "sorting score" was calculated using the cumulative GOF and cumulative LOF as shown in equation 9.

$$\text{cumulative GOF}_{(\mathrm{i})} = \int_{\mathrm{j}=1}^{\mathrm{i}} \text{norm GOF}_{(\mathrm{j})} \quad \text{for i} = 1,2..40 \ (7)$$

$$\text{cumulative LOF}_{(\mathrm{i})} = \int_{\mathrm{j}=1}^{\mathrm{i}} \text{norm LOF}_{(\mathrm{j})} \quad \text{for i} = 1,2..40 \ (8)$$

$$\text{sorting score}_{(\mathrm{i})} = \text{net GOF}_{(\mathrm{i})} - \text{net LOF}_{(\mathrm{i})} \ (9)$$

The clusters were then ranked based on those that contained the largest fraction of GOF relative to LOF mutations (Figure S3) as calculated by the cut-off score:

$$\text{cutoff score}_{(\mathrm{i})} = \text{cumulative GOF}_{(\mathrm{i})} - \text{cumulative LOF}_{(\mathrm{i})} \quad (10)$$

which revealed that most GOF or likely GOF (LGOF) mutations, and few LOF or likely LOF (LLOF) mutations, were in the top 10 clusters.

**Mutagenesis and plasmid preparation**

The following mutagenesis primers were used:

| Gene | Mutation | Forward Primer | Reverse Primer |
|---|---|---|---|
| PDGFRA | Y555C | AACAGAAACCGAGGTGTGAAATT CGCTGGAGGGTCAT | ATGACCCTCCAGCGAATTTCACA CCTCGGTTTCTGTT |
| | V561D | GAAATTCGCTGGAGGGACATTG AATCAATCAGCC | GGCTGATTGATTCAATGTCCCTC CAGCGAATTTC |
| | L580M | CTTGAGTCATAAGGCATCTGCAT CGGGTCCACA | TGTGGACCCGATGCAGATGCCT TATGACTCAAG |
| | Y849C | ACATCATGCATGATTCGA ACTGTGTGTCGA AAGGCAGTACCT TT | AAAGGTACTGCCTTTCGACACAC AGTTCGA ATCATGCATGATGT |
| | G652E | GAT AAT GAC TCA CCT GGA GCC ACA TTT GAA CAT TGT AA | TTA CAATGTTCA AATGTGGCTCCAGGTGAGTCATT A TC |
| FLT3 | Y572C | TGTAGCTGGCTTTCACACCTAAA TTGCTTTTTGTACTTGTGACA | TGTCACAAGTACAAAAAGCAATT TAGGTGTGAAAGCCAGCTACA |
| | M578I | GCCGGTCACCTGTACTATCTGTA GCTGGCTT | AAGCCAGCTACAGATAGTACAG GTGACCGGC |

| | | | |
|---|---|---|---|
| | F594I | GAGATCATATTCATATTCTCTGATATCAACGTAGAAGTACTCATTATCT | AGATAATGAGTACTTCTACGTTGATATCAGAGAATATGAATATGATCTC |
| | Y842C | GCCCCTGACAACACAGTTGGAATCACTCATGATATCTCG | CGAGATATCATGAGTGATTCCAACTGTGTTGTCAGGGGC |
| | G669E | CAATATTCTCGTGGCTTTCCAGCTGGGTCATCATC | GATGATGACCCAGCTGGAAAGCCACGAGAATATTG |
| | D835Y | GTTGGAATCACTCATGATATATCGAGCCAATCCAAAGTCAC | GTGACTTTGGATTGGCTCGATATATCATGAGTGATTCCAAC |
